# HTLV-1 reverse transcriptase homology model provides structural basis for sensitivity to existing nucleoside/nucleotide reverse transcriptase inhibitors

**DOI:** 10.1101/2023.05.25.542289

**Authors:** Nicolas Tardiota, Noushin Jaberolansar, Julia A. Lackenby, Keith J. Chappell, Jake S. O’Donnell

**Author notes:** **Corresponding author:** Jake S. O’Donnell.

## Abstract

The human T-lymphotropic virus type 1 (HTLV-1) infects millions of people globally and is endemic to various resource-limited regions. Infections persist for life and are associated with increased susceptibility to opportunistic infections and severe diseases including adult T cell leukemia/lymphoma (ATLL) and HTLV-1-associated myelopathy-tropical spastic paraparesis (HAM-TSP). No HTLV-1-specific anti-retrovirals have been developed and it is unclear whether existing anti-retrovirals developed for treatment of human immunodeficiency virus (HIV) have efficacy against HTLV-1. To understand the structural basis for therapeutic binding, homology modelling and machine learning were used to develop a structural model of the HTLV-1 reverse transcriptase. With this, molecular docking experiments using a panel of FDA-approved inhibitors of viral reverse transcriptases to assess their capacity for binding, and in turn, inhibition. Importantly, nucleoside/nucleotide reverse transcriptase inhibitor (NRTI) but not non-nucleoside reverse transcriptase inhibitors (NNRTIs) were capable of binding the HTLV-1 reverse transcriptase, with similar affinity to HIV-1 reverse transcriptase. By strengthening the rationale for clinical testing of therapies such as tenofovir alafenamide, zidovudine, lamivudine, and azvudine for treatment of HTLV-1, this study has demonstrated the power of *in silico* structural biology approaches in drug design and therapeutic testing.

## Introduction

Once established, human T-lymphotropic virus type 1 (HTLV-1) retroviral infections usually persist for life. While less severe than the closely related Human Immunodeficiency Virus (HIV), HTLV-1 infections result in sub-clinical immune suppression and are associated with a higher relative risk (RR) of all-cause mortality (RR 1.57; 95% CI 1.37–1.80) and a range of serious sequelae throughout life. Most seriously, HTLV-1 causes adult T cell leukemia/lymphoma (ATLL), a rare and extremely aggressive peripheral T cell cancer in 5% of cases, and HTLV-1-associated myelopathy-tropical spastic paraparesis (HAM-TSP), a degenerative autoimmune disease of the peripheral nervous system in a further 5% of cases [1-3]. Although uncommon in many developed countries, HTLV-1 is estimated to infect 10 to 20 million individuals globally [1-3].

No specific therapies have been developed to prevent, manage, or cure HTLV-1 infections, other than allogenic hematopoietic stem cell transplantation; a high-risk therapy used in treatment of aggressive ATLL [4]. Instead, interventions have been primarily limited to the management of HTLV-1-associated diseases, with limited success [1-3]. Adopting a pragmatic approach, research efforts have focused on testing anti-retroviral therapies developed for HIV against HTLV-1 such as Zidovudine (3’-azido-3’-deoxythymidine), tenofovir (9-(R)-[2-(phosphonomethoxy)propyl] adenine, PMPA), and lamivudine (2,3-dideoxy-3-thiacytidine) [5-7]. For many such compounds, *in vitro* testing has been able to demonstrate a successful reduction in proviral load [5-9]. However, of the few HTLV-1-related clinical studies performed, anti-retroviral therapies have not achieved this effect among chronically infected individuals [10]. One explanation for this discrepancy is that the HTLV-1 proviral load during chronic infection is maintained by reverse transcriptase-independent clonal proliferation [11]. By contrast, throughout the acute phase of infection, reverse transcriptase-mediated infective spread predominates and is critical for the establishment of a chronic infection [11]. This has led to the suggestion that anti-retroviral therapies might be more likely to suppress or eliminate an HTLV-1 infection when used as either pre- or post-exposure prophylaxis; approaches which have been particularly effective for HIV prevention and management [12]. Among at-risk populations, conducting clinicals trial to assess the effectiveness of pre- or post-exposure prophylaxis of reverse transcriptase inhibitors is possible; however, given that reporting of new HTLV-1 infections among adults is rare, the pool of patients available for inclusion in any trial is likely to be small. Therefore, more data are required to inform the rational selection of therapeutic candidates for inclusion.

To identify drug candidates likely to be of benefit, two outstanding questions must first be answered: (i) is structural similarity between HTLV-1 and HIV-1 sufficient to allow for binding of existing reverse transcriptase inhibitors? If so, (ii) do any of these inhibitors bind with sufficient affinity and in the correct conformation to inhibit HTLV-1 infective spread at a tolerable dose? The crystal structures of HTLV-1 retroviral proteins have not been resolved which has limited conventional structure-based analyses [13]. To bring greater attention to this neglected pathogen, we have addressed the above questions using *in silico* homology modelling and machine learning to predict a structural model of the HTLV-1 reverse transcriptase, something that has not previously been achieved. Using this model, we have performed molecular docking experiments to provide a framework to identify which, if any, existing retroviral reverse transcriptase inhibitors with FDA approval could be candidates for clinical testing against HTLV-1.

## Results

To address the above questions, it was first necessary to predict the structure of the HTLV-1 reverse transcriptase. Even among related species, DNA and amino acid sequences are often divergent. Despite this, protein structures tend to be highly conserved, presumably owing to the essential relationship between structure and function. By taking advantage of this, homology modelling can provide a theoretical prediction of a protein’s structure if the encoding DNA sequence is known and if crystal structure information is available for equivalent proteins of related species [14, 15]. The HIV-1 (protein databank identification number [PDBID]:1JLA), Moloney Murine Leukemia Virus (MMLV) (PDBID:4MH8), and Human Endogenous Retrovirus K (HERV-K) (PDBID:7SR6) retroviruses have been previously shown to share DNA and amino acid sequence similarity to HTLV-1 (NCIB: NC_001436) [16]. Based on previous annotations of HIV-1, MMLV, and HERV-K sequences, it was possible to infer within the HTLV-1 sequence, a 390 amino acid sequence (Gag-Pro-Pol amino acids 614 – 1004; annotated in NCIB:NC_001436) likely to contain all necessary domains to form the final reverse transcriptase structure. When comparing proteins, those with greater than 25% amino acid sequence similarity, usually take homologous 3D structures. Encouragingly, similarity was high between the identified HTLV-1 sequence and reverse transcriptases of HIV-1 (25%), MMLV (27%), and HERV-K (29%) (Figure 1) [14-16]. This provided confidence that the inferred amino acid sequence was highly likely to be associated with the HTLV-1 reverse transcriptase. To model the HTLV-1 reverse transcriptase structure, the identified 390 amino acid sequence was then input into Alphafold2; a machine learning algorithm which incorporates sequence homology, structural homology, secondary structure prediction, with contact maps (a ‘fingerprint’ of amino acid interactions in a folded structure), and has been reported to make highly accurate predictions of thousands of protein structures [17]. Through this process, Alphafold2 was able to generate 5 theoretical models of HTLV-1 Reverse transcriptase (Figure 2A and 2B). For further analyses, the model with the lowest predicted alignment error (PAE) score was used (Figure S1A). Further, this model demonstrated a typical resemblance to defined reverse transcriptases and had an obvious DNA binding pocket (Figure S1B). To determine the most energetically favourable conformation of the model and its proper molecular arrangement in 3D space, the structure was energy-minimized using GROMACS (Figure S1C).

**Figure 1.**
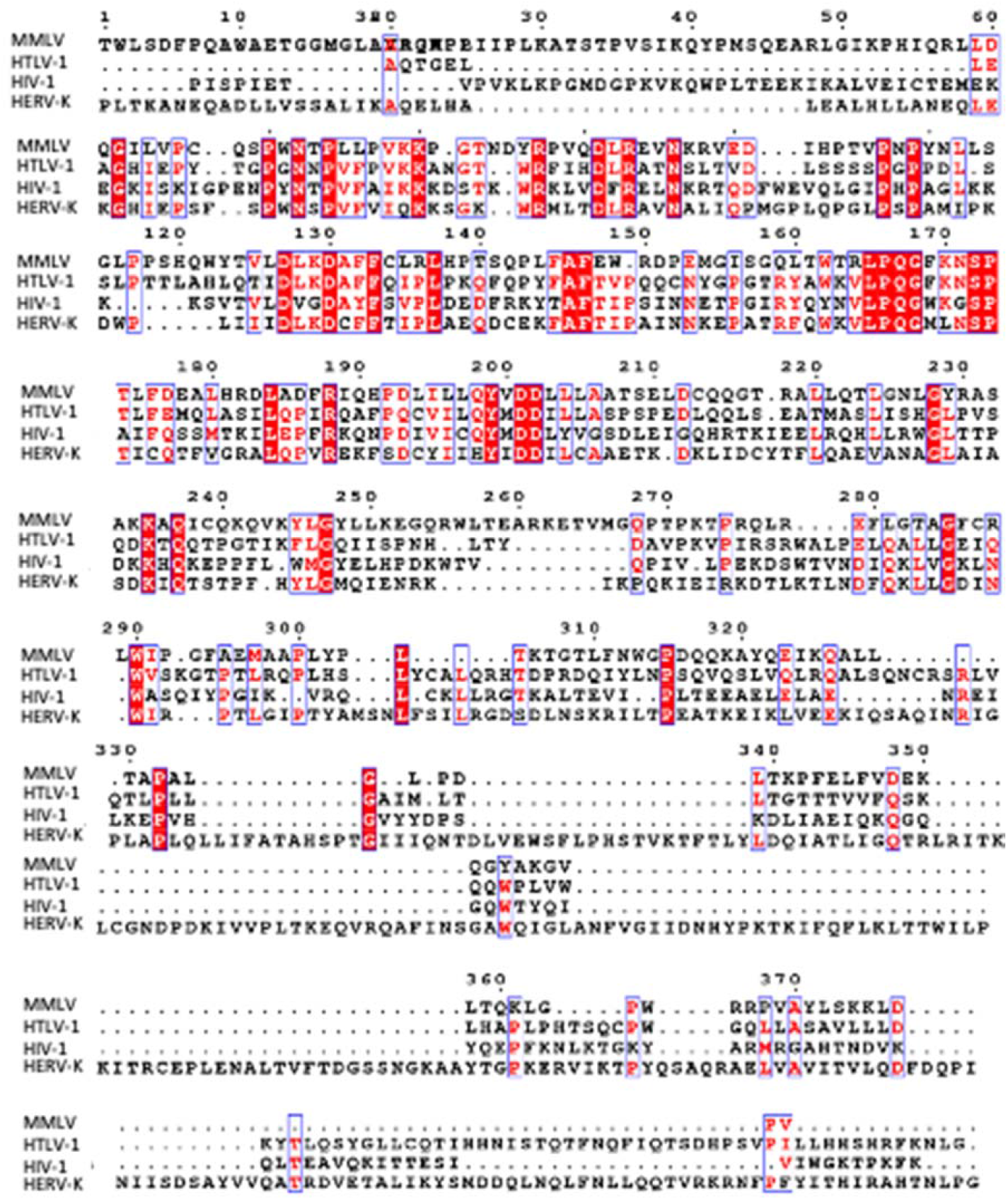
MMLV, HTLV-1, HIV-1, and HERV-K sequence alignment. Amino acid sequence alignment of MMLV, HTLV-1, HIV-1, and HERV-K. Amino acids conserved between all three viruses are highlighted in red with white text. For those residues considered for at least two viruses are outlined in blue with red text. Insertions are represented by a period.

**Figure 2.**
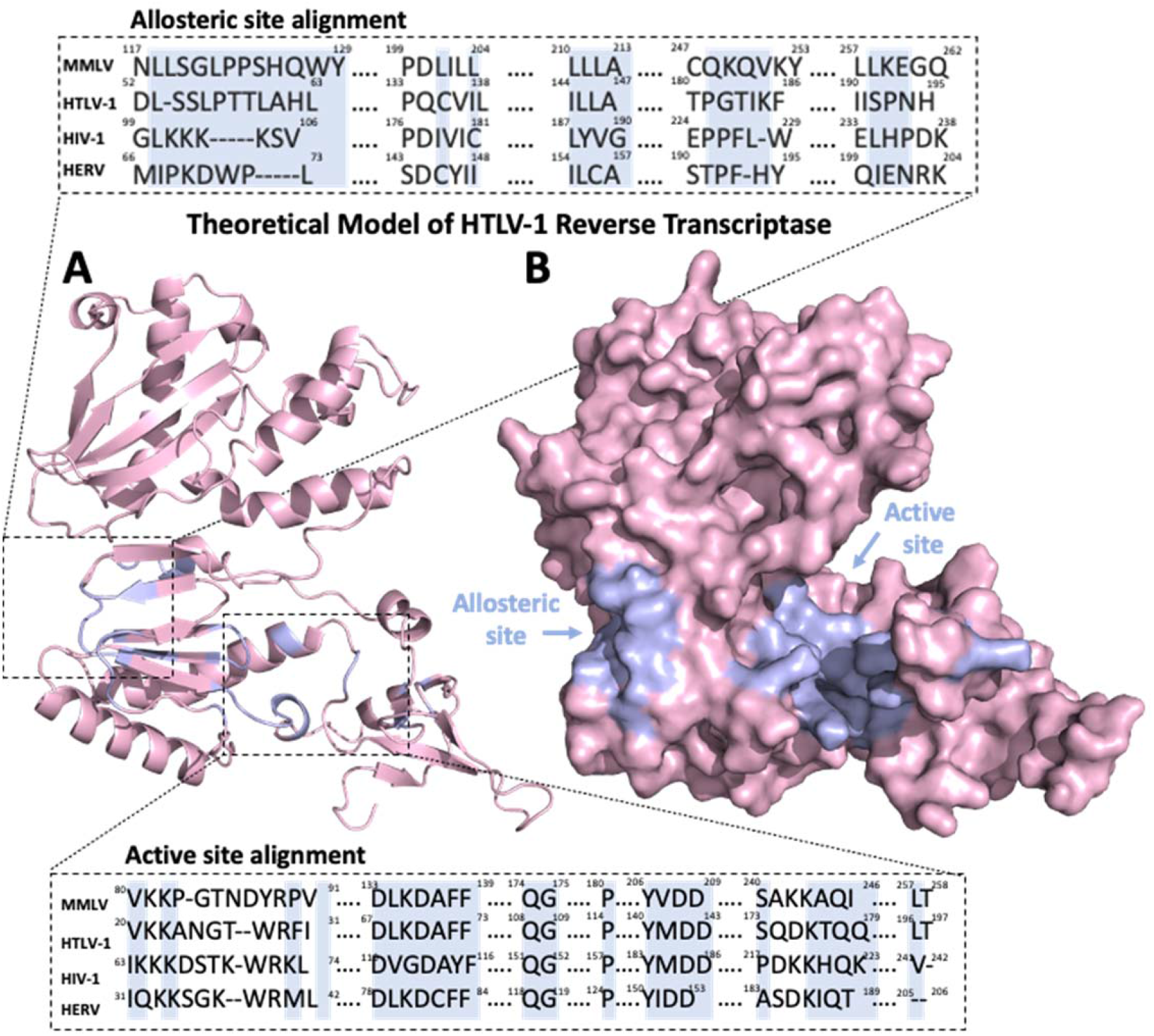
Rationalisation for theoretical model for HTLV-1 reverse transcriptase and molecular docking of reverse transcriptase inhibitors. **(A)** Cartoon representation of predicted HTLV-1 reverse transcriptase Alphafold2 model, and **(B)** its molecular surface. The predicted binding site of non-nucleoside reverse transcriptase inhibitors (NNRTIs) (allosteric site) and the binding site of nucleoside reverse transcriptase inhibitors (NRTIs) (active site) have been highlighted in purple. Amino acid sequence alignment for the active sites of HTLV-1 (NCIB: NC_001436), MMLV (PDBID:4MH8), HIV-1 (PDBID:1JLA), and HERV (PDBID:7SR6). Highlighted in purple are the amino acids defined for MMLV, HIV-1, and HERV (and predicted for HTLV-1) to interact with NNRTIs (above) and with NRTIs (below). **Related to Figure S1**.

To assess similarity between the predicted theoretical HTLV-1 model and those previously defined for MMLV (PDBID:4MH8), HERV-K (PDBID:7SR6), and HIV-1 (PDBID:1JLA), comparisons were made using root mean square deviation (R.M.S.D.); a commonly used quantitative measure of variation between superimposed atomic coordinates [18, 19]. Generally, R.M.S.D. values of <3.5 Å suggest a high degree of similarity (i.e. low structural variance). The HTLV-1 reverse transcriptase model was found to be highly structurally similar to MMLV (R.M.S.D. 0.109 Å); however, some structural variation was seen when it was compared to either HIV-1 (R.M.S.D. 4.282 Å) or HERV-K (R.M.S.D. 3.936 Å). It is worth noting; however, that these variations were modest in comparison to structural variation between HIV-1 and MMLV (R.M.S.D. 10.225 Å) (Figure 3A to 3B and Table 1).

**Figure 3.**
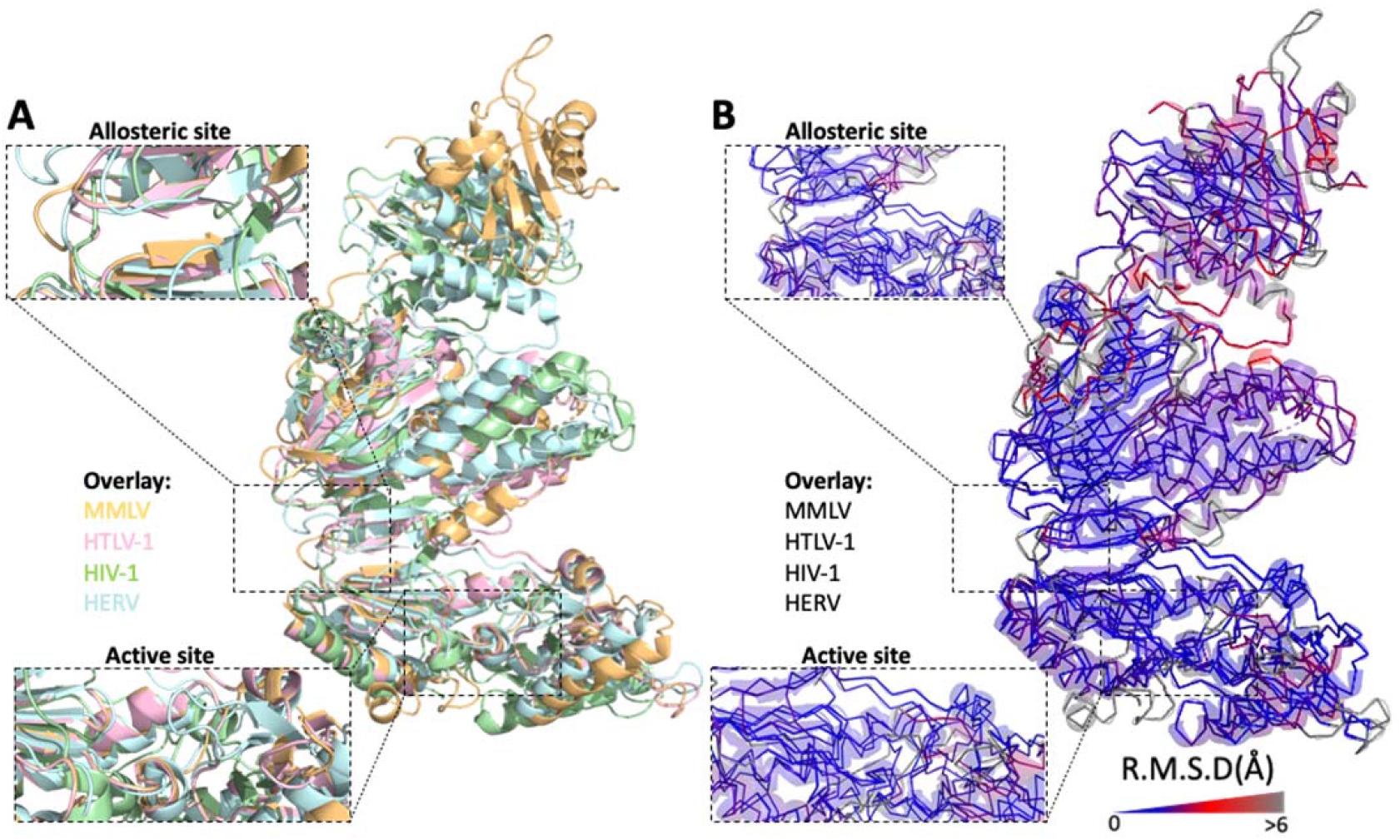
Reverse transcriptase structural comparison between MMLV, HTLV-1, HIV-1, and HERV-K. **(A)** Cartoon representation of HTLV-1 (pink), MMLV (orange), HERV (blue), and HIV-1 (green) reverse transcriptases (right). Inlays represent the active or allosteric site for each representation. **(B)** As for (A), ribbon diagram of backbone structural divergence measured as R.M.S.D. (Å) and depicted as blue (low) to grey (high) colour gradient (right). Inlays represent the active or allosteric site for each representation.

**Table 1.** Structural alignment of different species root mean square derivation score (R.M.S.D.) and homology model comparison for either whole reverse transcriptase structure, nucleoside reverse transcriptase inhibitor (NRTIs) binding site (active site), or non-nucleoside reverse transcriptase inhibitor (NNRTIs) binding site (allosteric site)

| Whole structure | 1jla | alpha | 4mh8 | 7sr6 |
| --- | --- | --- | --- | --- |
| 1jla | 100 |  |  |  |
| alpha | 4.282 | 100 |  |  |
| 4mh8 | 10.225 | 0.109 | 100 |  |
| 7sr6 | 3.956 | 3.936 | 11.075 | 100 |

| Allosteric | 1jla | alpha | 4mh8 | 7sr6 |
| --- | --- | --- | --- | --- |
| 1jla | 100 |  |  |  |
| alpha | 4.765 | 100 |  |  |
| 4mh8 | 1.847 | 0.083 | 100 |  |
| 7sr6 | 3.494 | 2.559 | 1.554 | 100 |

| Active | 1jla | alpha | 4mh8 | 7sr6 |
| --- | --- | --- | --- | --- |
| 1jla | 100 |  |  |  |
| alpha | 5.03 | 100 |  |  |
| 4mh8 | 2.563 | 1.3464 | 100 |  |
| 7sr6 | 4.746 | 4.332 | 4.244 | 100 |

| Whole structure | Phyre<sup>2</sup> | alpha | Swiss | Mod |
| --- | --- | --- | --- | --- |
| Phyre<sup>2</sup> | 100 |  |  |  |
| alpha | 3.109 | 100 |  |  |
| Swiss | 3.601 | 1.909 | 100 |  |
| Mod | 3.853 | 2.081 | 2.488 | 100 |

| Allosteric site | Phyre<sup>2</sup> | alpha | Swiss | Mod |
| --- | --- | --- | --- | --- |
| Phyre<sup>2</sup> | 100 |  |  |  |
| alpha | 1.109 | 100 |  |  |
| Swiss | 1.466 | 1.552 | 100 |  |
| Mod | 1.782 | 1.407 | 1.611 | 100 |

On the basis of previous annotations of HIV-1, it was possible to identify two sites within the HTLV-1 reverse transcriptase model likely to bind inhibitors of reverse transcriptase. These were the allosteric site which is targeted by non-nucleoside reverse transcriptase inhibitors (NNRTIs) and the active site which directly interacts with DNA and is the target of nucleoside analogue reverse transcriptase inhibitors (NARTIs or NRTIs) (Figure 2). Similar to the comparisons of the whole structures the HTLV-1 reverse transcriptase, when compared with MMLV, the active and allosteric sites showed minimal structural variation (R.M.S.D. allosteric site, 0.083 Å; active site, 1.346 Å). For comparisons between the HTLV-1 reverse transcriptase model active and allosteric sites and those of either HIV-1 (R.M.S.D. allosteric site, 4.765 Å; active site, 5.03 Å), or HERV-K (R.M.S.D. allosteric site, 2.559 Å; active site, 4.332 Å), structural variation was again seen (Figure 3A to 3B and Table 1). These findings suggest that despite sharing many structural characteristics, some overall and domain-specific structural variation exists between the HTLV-1 reverse transcriptase and those of HIV-1 and HERV-K, and to a lesser extent MMLV.

While advances in protein structural analysis are constantly being made, homology modelling can be an error-prone process, reliant on the assumptions and the data available to each software package. As such, orthogonal validation using crystal structure information is often important; however, when this information is unavailable, greater confidence in the predicted structure can be gained using separate methods [15, 20]. For example, using an identified amino acid sequence, it is possible to repeat structural modelling using separate software packages. From these separate models, structures identified to be generally similar are thought to be more likely representative of the protein’s native structure [15, 20]. For this, the inferred HTLV-1 reverse transcriptase amino acid sequence was input to three additional software packages: the Phyre^2^ protein folding web server [21], Modeller [22], and Swiss-Model, all of which build iterative models based on sequence and structural information (Figure 4A) [23]. Structural variation between the overall models was minimal (R.M.S.D. 1.909 Å - 3.853 Å), and further improved for the allosteric and active sites (R.M.S.D. 1.109 Å - 1.782 Å) (Table 1); providing greater confidence in the likelihood that the identified HTLV-1 amino acid sequence is representative of the native structure of the reverse transcriptase (Figure 4A to 4B) [18, 19]. Comparing the four predicted models, the greatest source of variation was introduced by the Modeller result, which by comparison, produced a model of HTLV-1 reverse transcriptase complexed with DNA, leading to larger shifts in secondary structure (Figure 4A to 4B).

**Figure 4.**
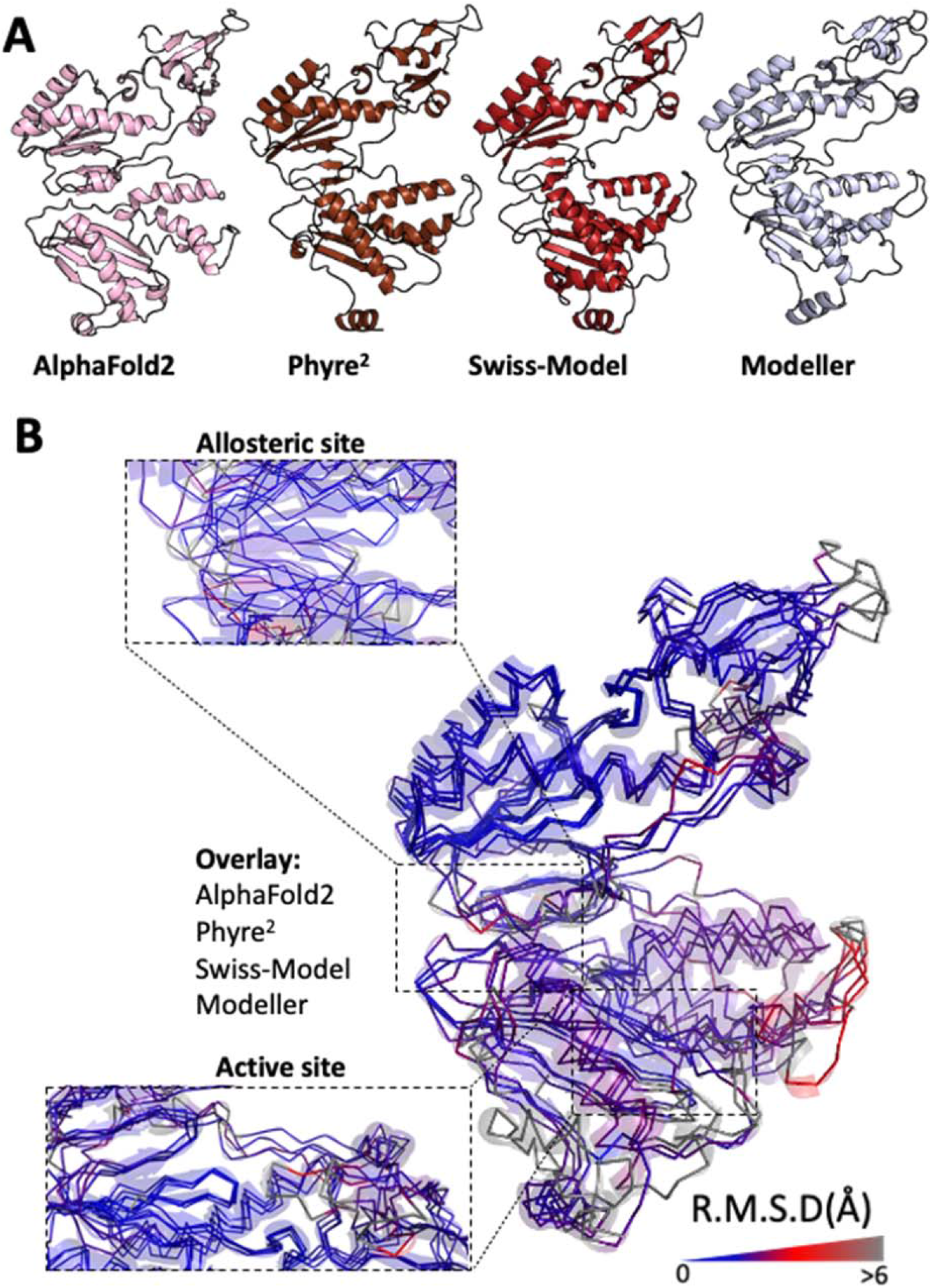
Modelling of theoretical HTLV-1 reverse transcriptase using alternative methods. **(A)** Cartoon representations of theoretical HTLV-1 reverse transcriptase modelled using Alphafold2, Phyre^2^, Swiss-Model, and Modeller. **(B)** As for (A), ribbon diagram of backbone structural divergence measured as R.M.S.D. (Å) and depicted as blue (low) to grey (high) colour gradient. Inlays represent the active or allosteric site for each representation.

Various *in vitro* studies, and a handful of clinical studies have suggested that some FDA-approved inhibitors of reverse transcriptase might have therapeutic activity against HTLV-1 [24]. As a cautionary note, as HTLV-1 is known to behave unusually *in vitro* meaning that it can be challenging to interpret these findings, and clinical studies performed to date have primarily focused on individuals with severe ATLL or HAM-TSP, which might confound results [24]. To provide greater clarity and context to these previous findings, we therefore wanted to test whether a structural basis for binding of these therapies exists within the HTLV-1 reverse transcriptase model, especially given that many of these therapies were designed to inhibit the HIV-1 reverse transcriptase allosteric and active sites. To do this, we selected four NNRTIs (rilpivirine, doravirine, nevirapine, and dapivirine) and four NARTIs or NRTIs (tenofovir alafenamide, zidovudine, lamivudine, and azvudine) to test with *in silico* docking experiments. These were performed using Autodock 4 which is capable of simulating interactions of molecules in different conformations within a protein structure and in doing so, can calculate interaction-associated binding energies (values <0 kcal/mol are favourable) [25]. Although other molecular docking programs exist, we chose Autodock 4 as it is able to handle molecule-ion interactions such as those which occur in the active site of the HTLV-1 reverse transcriptase model with Mg^2+^ (Figure S2D). As a control, we first tested molecular docking of each drug against the HIV-1 reverse transcriptase (PDBID:1JLA)(Figures S2A and S2C). NNRTIs were found to bind strongly in the allosteric site and each was tested in 10 different conformations providing binding energies ranging from -9.74 kcal/mol to -4.94 kcal/mol for doravirine, -7.02 kcal/mol to -1.49 kcal/mol for dapivirine, -8.27 kcal/mol to -4.39 kcal/mol for nevirapine, and -2.77 kcal/mol to 10.08 kcal/mol for rilpivirine (Figures S2A and S2C). Before molecular docking of NRTIs was attempted, they needed to be converted to biologically active, phosphorylated prodrug metabolites [26]. NRTIs were also found to bind strongly to the HIV-1 reverse transcriptase active site with -0.17 kcal/mol to 1.58 kcal/mol for tenofovir alafenamide, -1.43 kcal/mol to 0.71 kcal/mol for zidovudine, -1 kcal/mol to 1.55 kcal/mol for lamivudine, and -0.51 kcal/mol to 1.77 kcal/mol for azvudine (Figures 5A and 5C). In comparison with the HIV-1 reverse transcriptase allosteric site, the HTLV-1 reverse transcriptase allosteric site contained significantly more hydrophobic amino acids. Consequently, NNRTIs were unable to form hydrogen bonds in the allosteric site, yielding extremely poor binding affinities for 198.43 kcal/mol to 11.07 kcal/mol for doravirine, 75.58 kcal/mol to 6.69 kcal/mol for dapivirine, 27.15 kcal/mol to 15.38 kcal/mol for nevirapine, and 137.19 kcal/mol to 25.04 kcal/mol for rilpivirine (Figures S2B – S2C). This suggested that the NNRTIs tested might not have antiviral activity against HTLV-1. Surprisingly, despite R.M.S.D. differences between the HIV-1 and HTLV-1 reverse transcriptase active sites, interactions between NRTIs and the HTLV-1 reverse transcriptase active site were associated with improved binding energies, with -2.3 kcal/mol to 2.43 kcal/mol for tenofovir alafenamide, -2.94 kcal/mol to 0.71 kcal/mol for zidovudine, -2.26 kcal/mol to 1.43 kcal/mol for lamivudine, and -1.56 kcal/mol to 1.65 kcal/mol for azvudine (Figures 5A – 5C).

**Figure 5.**
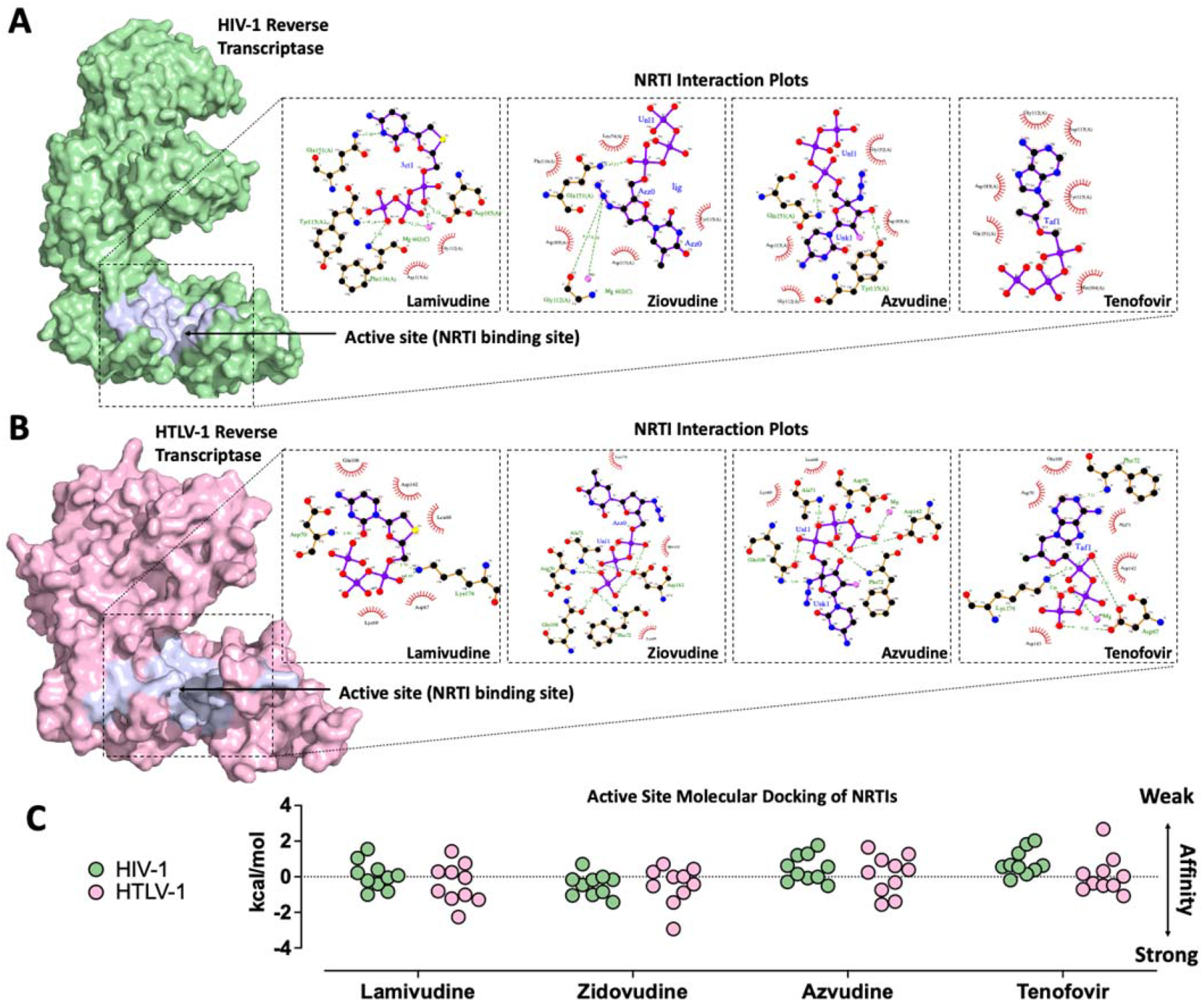
Molecular docking of reverse transcriptase inhibitors to HIV-1 and HTLV-1 reverse transcriptase. **(A)** Molecular surface diagram of HIV-1 reverse transcriptase with nucleoside reverse transcriptase inhibitor (NRTIs) binding site (active site) highlighted purple (left). Interaction plots of indicated NRTIs in the active site in their most energetically favourable conformation (1 of 10) (right). **(B)** Molecular surface diagram HTLV-1 reverse transcriptase with nucleoside reverse transcriptase inhibitor (NRTIs) binding site (active site) highlighted purple (left). Interaction plots of indicated NRTIs in the active site in their most energetically favourable conformation (1 of 10) (right). **(C)** Data summary of molecular docking testing 10 different conformations in either the HIV-1 reverse transcriptase or HTLV-1 reverse transcriptase. **Related to Figure S2**.

## Discussion

In this study we have used homology modelling and machine learning to develop a reasonable approximation of the HTLV-1 reverse transcriptase and used molecular docking to understand its binding interactions with FDA-approved inhibitors of reverse transcriptase. Together, these data suggest that chemical and structural dissimilarity between the reverse transcriptases of HIV-1 and HTLV-1 likely limits the efficient binding, and in turn potential for therapeutic efficacy of NNRTIs. Few if any studies have evaluated the capacity of NNRTIS such as rilpivirine, doravirine, nevirapine, and dapivirine to inhibit the HTLV-1 reverse transcriptase. Given their specificity for HIV-1, this is perhaps unsurprising. By contrast, the structural properties of the reverse transcriptase demonstrated clear and efficient binding to NRTIs which exceeded that of binding to the HIV-1 reverse transcriptase. These findings are important and support various preclinical and clinical studies performed to date. For example, zidovudine (AZT) has been demonstrated using *in vitro* and *in vivo* studies to be associated with decreased proviral load suggesting a capacity to limit infective spread [7, 27]. Although clinical studies investigating the role of AZT in treatment of HTLV-1 infection do not appear to have been performed, for treatment of HTLV-1-associated ATLL, AZT in combination with interferon α (IFNα) is currently recommended for treatment of symptomatic smouldering, unfavourable chronic, lymphoma (including extranodal primary cutaneous variant), and for acute disease with non-bulky tumor lesions [28]. From *in vitro* studies, it is clear that lamivudine has some capacity to protect lymphocytes from infection, however a methionine-to-valine substitution in the conserved motif of the HTLV-1 RT, tyrosine (Y)-methionine (M)-aspartic acid (D)-aspartic acid (D) (YMDD), has been shown to confer resistance in a similar way to how the M184V substitution which confers lamivudine resistance in HIV-1 RT [29]. In a small clinical study of patients with HAM-TSP lamivudine treatment coincided with a temporary decrease in circulating proviral load which rebounded back to baseline within 24 weeks of treatment [30]. Tenofovir has demonstrated *in vitro* inhibition of HTLV-1 reverse transcriptase; however, a small study in which daily treatment with 254 mg of tenofovir for a mean of 8.7 (+/- 2.3) months was not associated with a reduction in proviral load [31]. It is not clear whether azvudine has been tested for antiviral activity against HTLV-1.

Although clinical studies performed to date suggest that NRTIs have modest therapeutic benefit against HTLV-1, it is very important to recognise that the studies performed to date, have been on chronically infected individuals or those with severe ATLL or HAM/TSP. In chronically infected individuals, HTLV-1 viral activity is relatively quiescent, and reverse transcriptase-mediated infective spread contributes minimally to viral propagation, instead the proviral load is maintained by clonal proliferation. This suggests that targeting the HTLV-1 reverse transcriptase to treat chronically infected individuals might have limited efficacy [28]. By contrast, the acute phase which occurs in the months following infection is strongly associated with reverse transcriptase-mediated infective spread, meaning that this is the period during which an individual would be most likely sensitive to reverse transcriptase inhibition. This is the rationale for testing of these therapies using pre- and post-exposure prophylaxis regimens [28].

It is important to note that drugs targeting other retroviral proteins such as integrases and proteases do exist and are FDA-approved for various indications. A recent study used *in vitro* assays to identify HTLV-1 integrase inhibitors and found several candidates with potential activity against HTLV-1. Although these assays identified several drugs, the following *in silico* docking of these was only used to provide qualitative structural insight. For metal ion coordinated proteins such as integrases, it is currently difficult to use *in silico* approaches to derive quantitative molecular docking results as ion coordination presents challenges for existing software packages [32, 33].

HTLV-1 remains a neglected area of basic and clinical research. Following decades of intensive research on the pathogenesis of HIV, the tools now exist to understand the biology of HTLV-1 and for rational therapeutic development to take place. In this study, we aimed to understand whether a structural basis for binding to inhibitors of reverse transcriptase exists within the HTLV-1 reverse transcriptase. Limited by an unresolved protein structure, we developed and tested a theoretical model of HTLV-1 reverse transcriptase based on sequence alignment, homology modelling, and machine learning. Using this model, we identified that NRTIs such as tenofovir alafenamide, zidovudine, lamivudine, and azvudine are likely capable of binding and inhibition of the HTLV-1 reverse transcriptase.

## Methods

### Sequence alignment and homology modelling

To construct a viable sequence to use for de novo folding, homology modelling, and sequence alignment was done as a preliminary step to gauge an appropriate enzyme size with respect to number of amino acids. Using CLC Main Workbench (QIAGEN), the amino acid sequence of HTLV-1 Gag-Pro-Pol was aligned to that of HIV-1 reverse transcriptase (PDBID:1JLA), HERV-K reverse transcriptase (PDBID:7SR6), and MMLV reverse transcriptase (PDBID:4MH8) to look for conservation in sequence and infer the sequence for HTLV-1 reverse transcriptase. Using the pairwise analysis tool in CLC, a Point Accepted Mutation matrix (PAM) was constructed via the Dayhoff and Schwartz method (Dayhoff and Schwartz – Atlas of protein sequence and structure vol 3 of 5) to calculate the level of homology between proteins. In addition to sequence alignment was performed to enable, active site and allosteric site identification by structural alignment of HIV-1, HERV-K, and MMLV reverse transcriptases complexed with inhibitors where information was available: Allosteric site (PDBID; 1JLA, 1JLC, 1JEK),and Active site (PDBID; 5TXM, 7RS6, 4HKQ).

### De novo folding

The 390 amino acid sequence inferred to encode the HTLV-1 reverse transcriptase was input into the publicly available Alphafold2, Modeller, Swiss-Model, and Phyre^2^ web servers to produce structures as previously described [17, 18, 21, 23]. Related to the Modeller result, structural alignment was also performed with HERV (PDBID:7SR6).

### Energy minimisation

Energy minimisation was performed to relax the initial backbone conformation of the final reverse transcriptase structure and the active site. Using the GROMACS (5.31) simulation package a two-step energy minimization was done for a total of 2000 steps, with the first 1000 steps using the steepest decent method, followed by a further 1000 steps of conjugate gradient algorithm [34]. This was performed in explicit water using the tip4p water while interatomic interactions were modelled using AMBER force field (ffSB14) [35, 36].

### Molecular docking

The Alphafold2 structure was tested against a series of 8 different drugs in an *in silico* docking experiment, 4 NNRTIs in the allosteric site and 4 NRTI’s in the active site. These were carried out using the freely available AutoDock4 which is part of the Autodock Tools, which is a suite of programs used to prepare a protein and its corresponding drug target for Autodock4. These programs include Mgltools, PyMolecular View and Racoon [37]. Each drug target was localised to the predetermined interaction site, using the HIV-1 reverse transcriptase (PDBID:1JLA) and HERV-K (PDBID:7SR6) as a reference, with each docking experiment run through a series of 10 conformations via the generic search algorithm. The Mg^2+^ parameters were handled using an external parameter file, AD4_parameters.dat (https://autodock.scripps.edu/how-to-add-new-atom-types-to-the-autodock-force-field/).

### Ligand protein interactions

Visualisation of the ligands (NNRTI and NRTIs) and their associated interactions in the active and allosteric binding sites were visualised using Ligplot+. Using the best or lowest energy structure of the ligand in the binding pocket, interactions were visualised as either hydrogen bonds, dotted green lines or Van der Waals interactions, red semi-circles. Atoms were coloured using the CPK colouring method, while bonds were coloured purple. Confirmations with either more hydrogen bonds (green dotted lines) or more Van der Waals interactions, suggested a better fit (lower interaction energy and better ligand-protein interaction) at the binding site.

### Analysis

PyMol Molecular Graphics System Version 1.2r3pre (Schrödinger, LLC) was used to visualise folded protein structures and docking results. To compare target structures HIV-1, HERV-K, and MMLV to the HTLV-1 structure, the alignment tool was used and reported as the deviation of the backbone from the target structure (HIV-1) and reported in R.M.S.D. in Å. Blue to red scale was used to represent an approximation of backbone deviation using the colourbyrmsd plug-in.

## Author Contributions

Study Conception: NT, JSOD

Computational modelling and analysis: NT, NJ, JAL, JSOD

Writing: JSOD, NT

Editing: NT, NJ, KJC, JAL

Supervision: JSOD, KJC

## Acknowledgements and Conflicts of Interest

Keith Chappell declares holdings in ViceBio Limited and has patents pending related to molecular clamp platform (AU 2018241252; BR112019019813.9; CA 3057171; CH 201880022016.9; EP 18775234.0; IN 201917038666; ID P00201909145; IL 269534; JP 2019-553883; MX/a/2019/011599; NZ 757178; KR 0-2019-7031415; SG 11201908280 S; US 16/498865). All remaining authors declare no conflicts of interest.

## References

1. Yoshida, M., I. Miyoshi, and Y. Hinuma, Isolation and characterization of retrovirus from cell lines of human adult T-cell leukemia and its implication in the disease. Proc Natl Acad Sci U S A, 1982. 79(6): p. 2031–5.

2. Schierhout, G., et al., Association between HTLV-1 infection and adverse health outcomes: a systematic review and meta-analysis of epidemiological studies. Lancet Infect Dis, 2020. 20(1): p. 133–143.

3. Koga, Y., et al., Trends in HTLV-1 prevalence and incidence of adult T-cell leukemia/lymphoma in Nagasaki, Japan. J Med Virol, 2010. 82(4): p. 668–74.

4. Katsuya, H., et al., Treatment and survival among 1594 patients with ATL. Blood, 2015. 126(24): p. 2570–7.

5. Taylor, G.P., et al., Effect of lamivudine on human T-cell leukemia virus type 1 (HTLV-1) DNA copy number, T-cell phenotype, and anti-tax cytotoxic T-cell frequency in patients with HTLV-1-associated myelopathy. J Virol, 1999. 73(12): p. 10289–95.

6. Hill, S.A., et al., Susceptibility of human T cell leukemia virus type I to nucleoside reverse transcriptase inhibitors. J Infect Dis, 2003. 188(3): p. 424–7.

7. Matsushita, S., et al., Pharmacological inhibition of in vitro infectivity of human T lymphotropic virus type I. J Clin Invest, 1987. 80(2): p. 394–400.

8. Marino-Merlo, F., et al., Antiretroviral Therapy in HTLV-1 Infection: An Updated Overview. Pathogens (Basel, Switzerland), 2020. 9(5): p. 342.

9. Izaki, M., et al., In vivo dynamics and adaptation of HTLV-1-infected clones under different clinical conditions. PLoS Pathog, 2021. 17(2): p. e1009271.

10. Treviño, A., et al., Antiviral effect of raltegravir on HTLV-1 carriers. Journal of Antimicrobial Chemotherapy, 2011. 67(1): p. 218–221.

11. O’Donnell, J.S., Hunt Stewart K., Chappell Keith J., Integrated molecular and immunological features of HTLV-1 infection and disease progression to adult T cell leukemia/lymphoma. Lancet Haematology, 2023.

12. Bradshaw, D. and G.P. Taylor, HTLV-1 Transmission and HIV Pre-exposure Prophylaxis: A Scoping Review. Front Med (Lausanne), 2022. 9: p. 881547.

13. Sarafianos, S.G., et al., Crystal structure of HIV-1 reverse transcriptase in complex with a polypurine tract RNA:DNA. Embo j, 2001. 20(6): p. 1449–61.

14. Rost, B. and C. Sander, Bridging the protein sequence-structure gap by structure predictions. Annu Rev Biophys Biomol Struct, 1996. 25: p. 113–36.

15. Muhammed, M.T. and E. Aki-Yalcin, Homology modeling in drug discovery: Overview, current applications, and future perspectives. Chem Biol Drug Des, 2019. 93(1): p. 12–20.

16. Soltani, A., et al., Molecular targeting for treatment of human T-lymphotropic virus type 1 infection. Biomedicine & Pharmacotherapy, 2019. 109: p. 770–778.

17. Jumper, J., et al., Highly accurate protein structure prediction with AlphaFold. Nature, 2021. 596(7873): p. 583–589.

18. Wallner, B. and A. Elofsson, All are not equal: a benchmark of different homology modeling programs. Protein Sci, 2005. 14(5): p. 1315–27.

19. Haddad, Y., V. Adam, and Z. Heger, Ten quick tips for homology modeling of high-resolution protein 3D structures. PLoS Comput Biol, 2020. 16(4): p. e1007449.

20. Nikolaev, D.M., et al., A Comparative Study of Modern Homology Modeling Algorithms for Rhodopsin Structure Prediction. ACS Omega, 2018. 3(7): p. 7555–7566.

21. Kelley, L.A., et al., The Phyre2 web portal for protein modeling, prediction and analysis. Nature Protocols, 2015. 10(6): p. 845–858.

22. Martí-Renom, M.A., et al., Comparative protein structure modeling of genes and genomes. Annu Rev Biophys Biomol Struct, 2000. 29: p. 291–325.

23. Waterhouse, A., et al., SWISS-MODEL: homology modelling of protein structures and complexes. Nucleic Acids Res, 2018. 46(W1): p. W296–w303.

24. Marino-Merlo, F., et al., Antiretroviral Therapy in HTLV-1 Infection: An Updated Overview. Pathogens, 2020. 9(5).

25. Forli, S., et al., Computational protein–ligand docking and virtual drug screening with the AutoDock suite. Nature Protocols, 2016. 11(5): p. 905–919.

26. De Clercq, E. and H.J. Field, Antiviral prodrugs - the development of successful prodrug strategies for antiviral chemotherapy. Br J Pharmacol, 2006. 147(1): p. 1–11.

27. Isono, T., K. Ogawa, and A. Seto, Antiviral effect of zidovudine in the experimental model of adult T cell leukemia in rabbits. Leuk Res, 1990. 14(10): p. 841–7.

28. O’Donnell, J.S., Hunt, S.K., Chappell, K.J., Integrated molecular and immunological features of HTLV-1 infection and disease progression to adult T cell leukemia/lymphoma The Lancet Haematology 2023. Under Review.

29. García-Lerma, J.G., S. Nidtha, and W. Heneine, Susceptibility of human T cell leukemia virus type 1 to reverse-transcriptase inhibitors: evidence for resistance to lamivudine. J Infect Dis, 2001. 184(4): p. 507–10.

30. Taylor, G.P., et al., Effect of Lamivudine on Human T-Cell Leukemia Virus Type 1 (HTLV-1) DNA Copy Number, T-Cell Phenotype, and Anti-Tax Cytotoxic T-Cell Frequency in Patients with HTLV-1-Associated Myelopathy. Journal of Virology, 1999. 73(12): p. 10289–10295.

31. Macchi, B., et al., Susceptibility of primary HTLV-1 isolates from patients with HTLV-1-associated myelopathy to reverse transcriptase inhibitors. Viruses, 2011. 3(5): p. 469–83.

32. Barski, M.S., et al., Structural basis for the inhibition of HTLV-1 integration inferred from cryo-EM deltaretroviral intasome structures. Nature Communications, 2021. 12(1): p. 4996.

33. Li, P. and K.M. Merz, Metal Ion Modeling Using Classical Mechanics. Chemical Reviews, 2017. 117(3): p. 1564–1686.

34. Berendsen, H.J.C., D. van der Spoel, and R. van Drunen, GROMACS: A message-passing parallel molecular dynamics implementation. Computer Physics Communications, 1995. 91(1): p. 43–56.

35. Abascal, J.L.F. and C. Vega, A general purpose model for the condensed phases of water: TIP4P/2005. The Journal of Chemical Physics, 2005. 123(23): p. 234505.

36. Maier, J.A., et al., ff14SB: Improving the Accuracy of Protein Side Chain and Backbone Parameters from ff99SB. J Chem Theory Comput, 2015. 11(8): p. 3696–713.

37. Morris, G.M., et al., AutoDock4 and AutoDockTools4: Automated docking with selective receptor flexibility. J Comput Chem, 2009. 30(16): p. 2785–91.

